# PathwayPilot: A User-Friendly Tool for Visualizing and Navigating Metabolic Pathways

**DOI:** 10.1101/2024.06.21.599989

**Authors:** Tibo Vande Moortele, Pieter Verschaffelt, Qingyao Huang, Nadezhda T. Doncheva, Tanja Holstein, Caroline Jachmann, Peter Dawyndt, Lennart Martens, Bart Mesuere, Tim Van Den Bossche

## Abstract

**Background:** Metaproteomics, the study of collective proteomes in environmental communities, plays a crucial role in understanding microbial functionalities affecting ecosystems and human health. Pathway analysis offers structured insights into the biochemical processes within these communities. However, no existing tool effectively combines pathway analysis with peptide- or protein-level data.

**Results:** This manuscript introduces PathwayPilot, a user-friendly web application for exploring and visualizing metabolic pathways. PathwayPilot can compare functional annotations across different samples or organisms within a sample. A case study on the impact of caloric restriction on gut microbiota demonstrated the tool’s efficacy in deciphering complex metaproteomic data. The re-analysis revealed significant shifts in enzyme expressions related to short-chain fatty acid biosynthesis, aligning with existing research findings and showcasing PathwayPilot’s capability for accurate functional annotation and comparison across different microbial communities.

**Conclusions:** PathwayPilot represents a significant advancement in metaproteomic data analysis, offering a user-friendly interface for exploring and visualizing metabolic pathways. This study not only validates the tool’s applicability in real-world scenarios but also highlights its potential for broader research implications in microbial ecology and health sciences.

## Background

Metaproteomics, the study of the collective proteome of environmental communities, has emerged as a powerful tool to study microbiomes. These communities, often composed of a myriad of diverse microorganisms, play pivotal roles in shaping ecosystems, biogeochemical cycles, and human health. Understanding the functional capabilities of these diverse microbial communities is therefore essential for unraveling their ecological roles, potential biotechnological applications, and contributions to various ecosystem processes. While nucleotide-based methods such as metagenomics are often used to study microbiomes, metaproteomics is able to identify the temporal and spatial abundance of metabolic enzymes and other proteins. Metaproteomics is therefore complementary to nucleotide-based methods to learn about the real-time, functional state of the microbiome [1].

There are several methods to study functions in microbiomes, including but not limited to comparing Gene Ontology (GO) terms [2–4], Enzyme Commission (EC) numbers [5], and visualizing metabolic pathways, particularly those cataloged in the Kyoto Encyclopedia of Genes and Genomes (KEGG) database [6]. These pathways provide a structured framework to encapsulate the biochemical processes and functions involved in a microbial community. KEGG pathways, in particular, serve as an extensive repository of curated knowledge about these cellular processes, offering a way to decipher the functional landscape of the microbiome. Therefore, they help to address critical questions, such as “How do microbial communities respond to environmental changes?”, “What functional roles do specific microorganisms play within a community?”, and “How do these roles shift in response to perturbations or ecological shifts?”. Pathway analysis therefore offers valuable insights into metabolic, signaling, and regulatory networks present in the microbial community.

Currently, there is no tool that allows combining pathways with peptide-level or protein-level information in order to help answering these questions. Even more so, when we look at multiple species and conditions. It is therefore important to provide new user-friendly tools for these analyses to propel the field forward [7].

In this manuscript we introduce PathwayPilot, a versatile tool designed to tackle these challenges. This user-friendly web application is freely available at https://pathwaypilot.ugent.be. Its source code is available under the MIT open-course license at https://github.com/unipept/pathway-pilot. Specifically, PathwayPilot is designed to explore and visualize metabolic pathways. It has the capability to (quantitatively) compare functional annotations associated with a single organism across different samples and also to compare functional annotations across multiple organisms within a single sample. It accepts both peptides and proteins as input, allowing users to choose between peptide-centric and protein-centric analysis for metaproteomics, as both options are still often used in the field [8]. By mapping identified peptides or proteins onto EC numbers, taxon identifiers, and KEGG metabolic pathway maps, PathwayPilot can provide a clear and intuitive visualization of metaproteomics data. With our user-friendly interface, users can easily navigate through the pathways and highlight specific peptides, proteins or species of interest.

## Implementation

### PathwayPilot’s architecture

PathwayPilot is an interactive web application. Its frontend is written in TypeScript and based on a combination of Vue 3 and Vuetify. Vue is a powerful web framework that enables the implementation of high-performance, interactive applications by providing a layer of reactivity that enhances visualization updates. Vuetify, on the other hand, streamlines the development process by handling the intricacies of design and the creation of complex data tables, buttons, and other UI elements.

The backend is written in TypeScript, coupled with NodeJS. NodeJS’s V8 JavaScript engine stands out for its exceptional efficiency and performance, aligning perfectly with PathwayPilot’s requirements, and it shares its programming language with the frontend, thus simplifying code maintenance.

PathwayPilot relies on a dedicated backend server, which optimizes the entire mapping process from functional information to pathways and solves some browser and HTTP header limitations. All data is thus centralized at the server and can be requested dynamically by the frontend to provide a better user experience, and better analyses opportunities. The entire pipeline process is illustrated in **Figure 1**.

**Figure 1:**
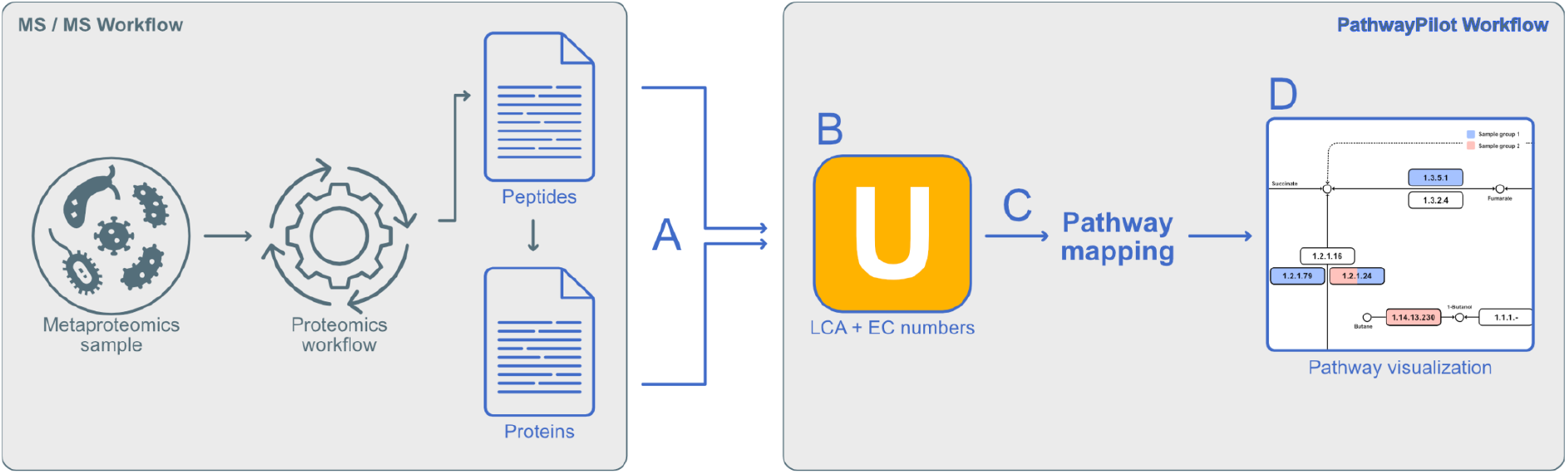
PathwayPilot workflow from sample to visualization. Gray arrows indicate the process prior to the pathway analysis. The blue arrows are part of the PathwayPilot processing stages. No user interaction is needed for these internal parts. A) User uploads either peptides or proteins. B) Unipept’s API is used to annotate the input data. C) The input peptides or proteins are annotated with pathway information. D) The annotated data is presented to the user.

### Supporting peptide and protein inputs

PathwayPilot supports both peptide and protein based input formats. However, dealing with different formats presented two main challenges.

First, upon data upload, all of the peptide/protein, functional, taxonomic, and pathway mappings are generated and interlinked. This layer then allows user interactions to be seamless and fluid, as it obviates the need for extensive recalculations at each interaction.

Second, in order to map two different input formats onto our shared internal representation, we ideally need two endpoints with similar outputs: one for peptides, and another for proteins (represented as UniProtKB accession numbers). To retrieve peptide-based taxonomic and functional information, the existing Unipept *peptinfo* endpoint can be used. This endpoint can handle large peptide batches at once, and provides results in an easy-to-use JSON format. For proteins, Unipept [9] provides a novel *protinfo* endpoint (release 5.0.9). This new, protein-focused endpoint mirrors the output format of the *peptinfo* endpoint, and can also handle batches of proteins rather than individual protein requests. This enhancement drastically improves data analysis speed as compared to using the UniProtKB endpoint, which only supports single requests, or the UniProtKB’s [10] XML format, requiring additional processing.

### Mapping peptides and proteins to KEGG pathways

KEGG pathways are a collection of directed cyclic graphs with labeled nodes and edges. They represent knowledge of molecular interactions, reactions and relations. Kegg pathways consist of four distinct and interconnected graph elements, as displayed in **Figure 2**. The first element represents compounds, identified by the ‘C’ prefix. The corresponding node is represented by a circle, except when shown in overview maps. The second element represents functions, and is characterized by KEGG ontology identifiers (K) and Enzyme Commission numbers (EC). As nodes, these are shown as rectangular labels on top of edges. The third element represents reactions, identified by the ‘R’ prefix and represented by (bi-)directional solid lines and arrows. The fourth and final element is used to interconnect different pathways, and these pathway links are shown as dashed lines.

**Figure 2:**
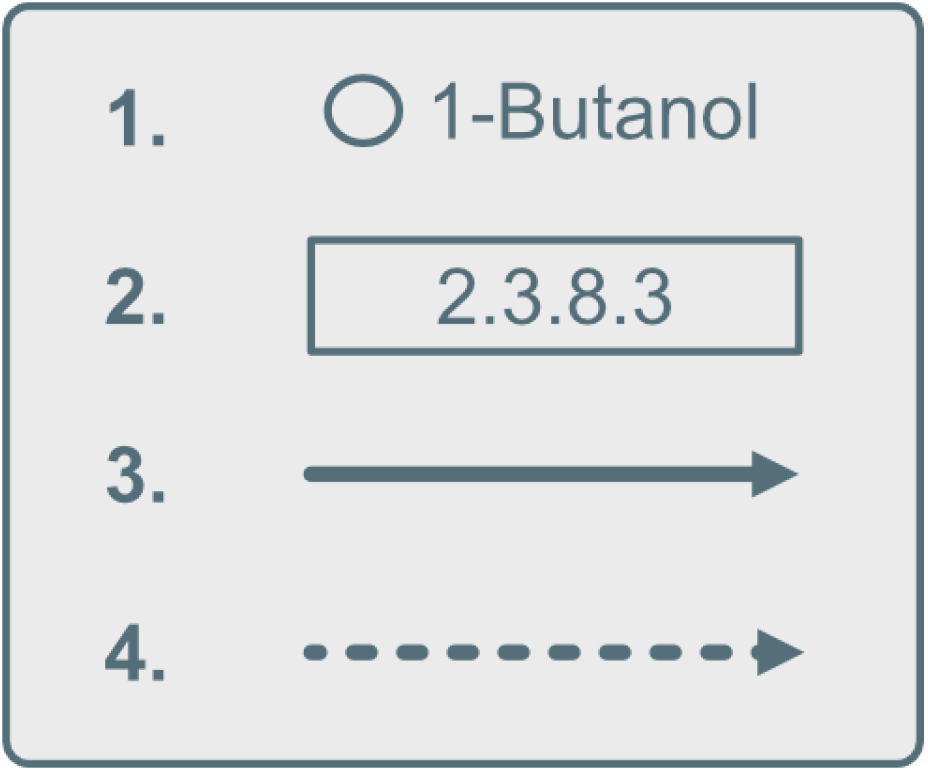
Different KEGG pathway elements. 1) Compound nodes, mostly accompanied by their name. 2) Enzymes used by a reaction. The area mostly contains an EC number linked to the enzyme. 3) Reaction edges, indicating a reaction between compounds. 4) Pathway edges, interconnecting two neighboring pathways.

PathwayPilot exclusively works with EC numbers, so in order to link peptides and proteins to KEGG pathways, we must first adapt all pathway node information to align with these EC numbers. This adaption includes all compounds, reactions, functions and pathway information. This information is retrieved from link files containing all essential mappings, enabling us to query node information using an EC number as the reference point.

Additionally, we need to map user-provided peptides and UniProtKB accession numbers to EC numbers. For the mapping of tryptic peptides, we use the Unipept API [11]. Unipept performs an *in silico* tryptic peptide digest of the entire UniProtKB database in its construction, facilitating rapid searches based on its built-in peptide index. For proteins, the novel *protinfo* endpoint leverages the fast Unipept API to provide a direct link from protein to EC number. Note that consolidating this approach around Unipept brings the advantage of uniform data sources across both input formats, as both rely on the same data source.

We thus end up with two separate mappings: one connecting peptides or proteins to EC numbers, and another that links EC numbers to pathway information. The mapping of peptides or proteins to a selected pathway is then transitively obtained by matching the EC numbers within the dataset to the calculated annotations found in pathway nodes. To optimize pathway data management, we have implemented a reverse proxy to serve as an intermediary between KEGG and PathwayPilot. This proxy simplifies the data retrieval process by directly downloading the pathway’s webpage, extracting the relevant image and node information, and performing essential preprocessing steps. By doing so, it drastically reduces the number of API requests needed, thus improving performance and reducing the complexity of client-side operations.

### Comparative data visualization

PathwayPilot provides two types of visualization. First, there is the standard visualization, which highlights nodes with at least one match between user input and a pathway node. Second, we developed a visualization that shows the normalized difference between two groups. In both cases, the KEGG pathway image is retrieved from the proxy server, and a transparent vector graphic (SVG) is constructed to overlay the image.

To visualize matches within pathways, a simple comparison between the uploaded data and each pathway node’s information is conducted. Each match is then coloured in the SVG overlay, superimposed on the KEGG pathway image. When multiple groups are visualized on a pathway, the overlay nodes are segmented into multiple parts, each assigned a separate color.

To visualize the difference in abundance between two groups of peptides, based on the absolute number of matches within a pathway node, we calculate the difference as depicted in **Formula 1**.

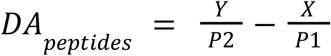

**Formula 1**: Calculation of the differential abundance of peptides. X represents the number of matched peptides in the first group and Y in the second group. P1 and P2 represent the total number of peptides in the first and second group. By using the total number of peptides as a denominator, we prevent a bias towards the bigger group in terms of peptides. Positive values indicate a higher abundance in the first group, while negative values indicate more matches in the second group.

For proteins, we can’t use an absolute protein count to visualize the abundance between multiple groups. However, because we accept an abundance number alongside the protein accession number, we can use the *log*_2_ *fold change*. This is an often used metric to compare regulation between two different sample groups. We can then calculate the difference as depicted in **Formula 2**.

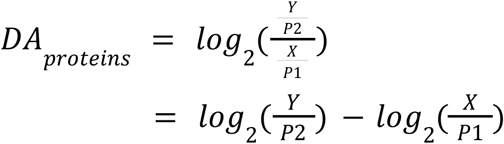

**Formula 2**: Calculation of the differential abundance of proteins. X and Y represent the sum of protein abundances in the first and second group respectively. P1 and P2 represent the total number of proteins in the first and second group. A log_2_ fold-change is calculated to define the difference between the two groups.

However, coloring nodes based on these differences could result in misleading visualizations because the color range is defined by the minimum and maximum values. If either the absolute minimum or maximum is significantly smaller or bigger than the other, the color may inaccurately shift towards one of the groups. To address this, we normalize the values on both ends and use a diverging scale to fix the midpoint, effectively combining two sequential scales around a critical midpoint.

## Results

The PathwayPilot interface consists of four distinct steps (**Figure 3 - 5**). First, users upload one or multiple datasets. Next, they select the pathway they wish to analyze, with tools in place to ease this process. Subsequently, users can analyze the pathway interactively using an image. Finally, they can create custom downstream analyses using various export options. To ensure efficiency and user-friendliness, we developed PathwayPilot as a single-page application that minimizes clicks by breaking the analysis process into a few concise and logical steps. All action elements are easy to find and serve a single function, adhering to Material Design guidelines [12] for intuitive use.

**Figure 3:**
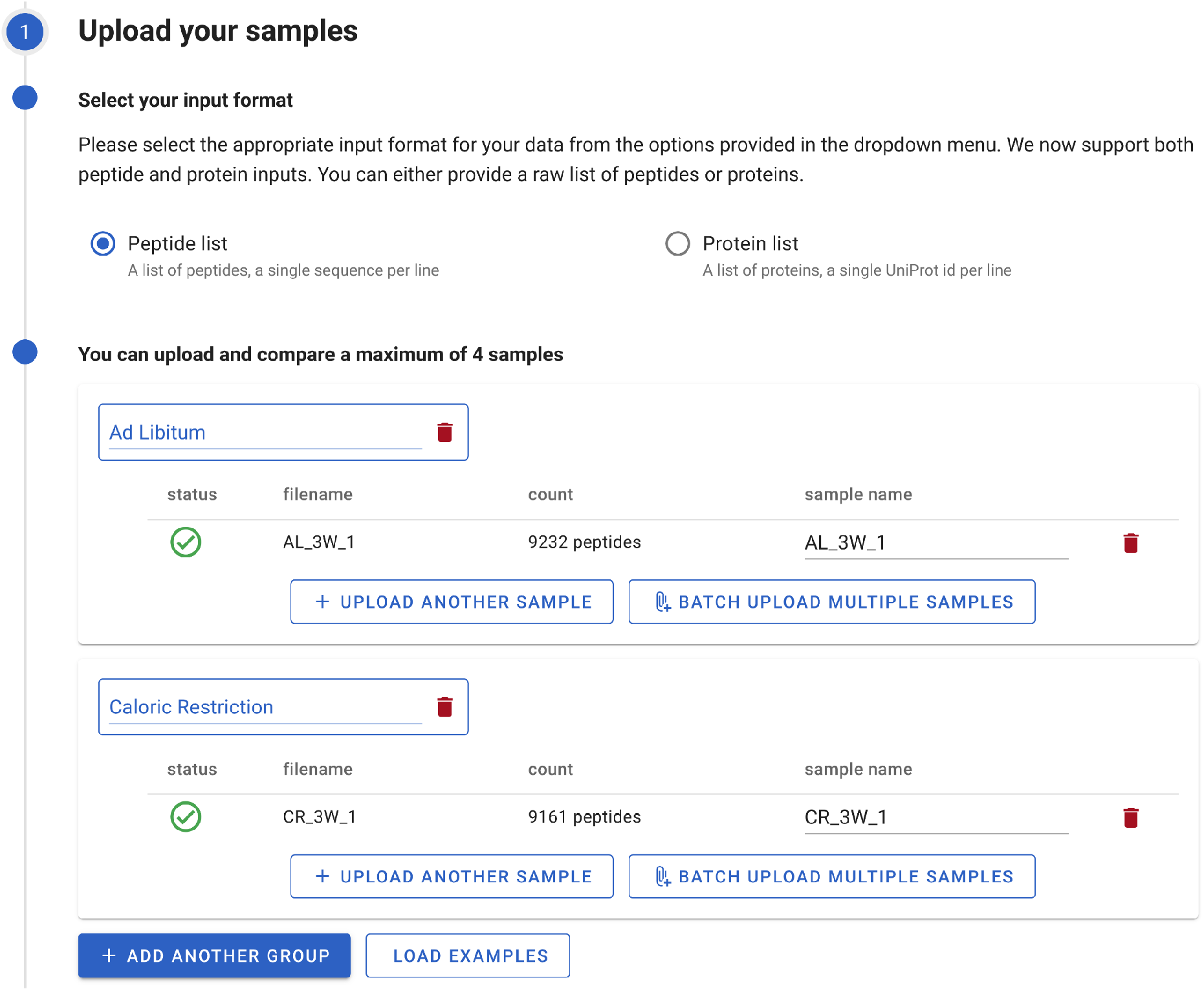
Data uploading interface. The user chooses between peptides or proteins as input data, and uploads either a single sample, a group of samples, or multiple groups of samples to compare.

### 1. Uploading the samples

After choosing the type of analysis, either to compare organisms or to compare a few different sample groups, data upload can start. The uploading process is visualized in **Figure 3**.

First, select the format of the data. Because each format behaves slightly differently in the subsequent steps, it is important to commit to either peptide- or protein-based analysis. During any analysis, all uploaded files have to conform to the same format. Changing the format during successive steps in the analysis will delete all data and restart the analysis.

Second, upload the data. This step is highly similar for both types of analysis. The only differences are the optional division in groups and the batched import when working with more than one sample. In case the user only wants to compare organisms or a single sample, they can paste their sequences or upload a single file using the file selector. Upon upload, PathwayPilot first checks whether the file is valid, and whether it conforms to the selected input format. If the file is found to be invalid, PathwayPilot will notify the user before starting the effective upload. This ensures that a user will not have to wait for the upload of a file only to have that upload fail at the end, due to, say, a formatting error in the last line of the file. Each malformed line detected will be reported, alongside with the line number and the reason.

### 2. Selecting a pathway

In the next step, choose a pathway for visualization (**Figure 4)**. All pathways are shown in order of matches against the data uploaded in the previous step, i.e. the number of peptides or proteins mapped to the specific pathway. This count aggregates matches across all uploaded samples. PathwayPilot offers two options to select a pathway. Either select a row from a sorted table or select a bubble from the provided bubble plot. Both interact seamlessly, ensuring a uniform and straightforward selection process.

**Figure 4:**
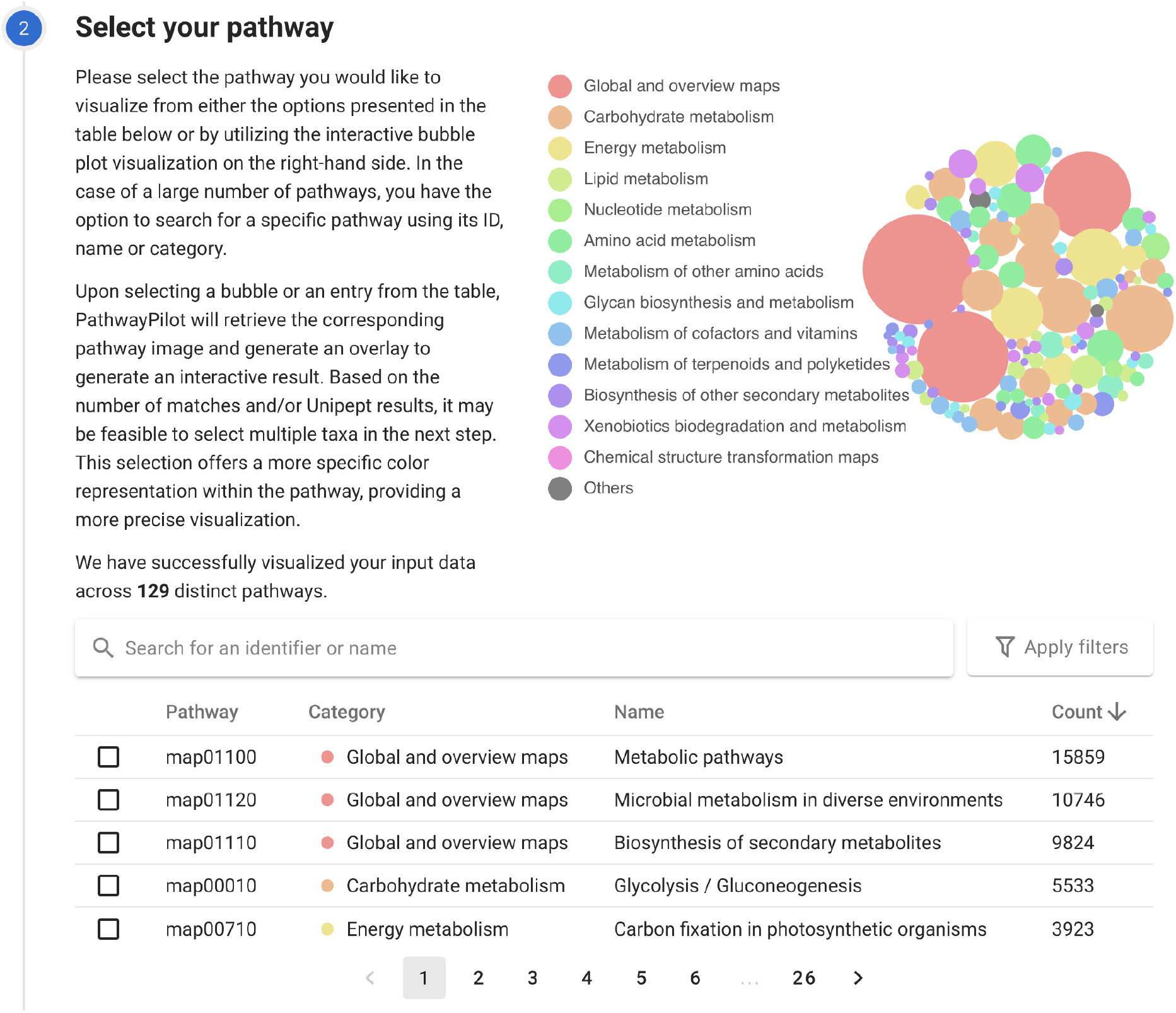
Pathway selection. The user selects a pathway, and can use the search bar or advanced filters to quickly find a specific pathway.

Additionally, two filtering options were added. The search bar allows textual filters based on pathway categories, identifiers or names. Separate filtering on the other hand allows inclusion of pathways that contain specific enzymes (EC numbers) or compounds (C numbers). These filtering options can all be combined to construct more specific queries. PathwayPilot’s preprocessing ensures that only the filter options leading to at least one result are shown.

### 3. Analyzing the pathway

Upon uploading the data and selecting the pathway for visualization, PathwayPilot starts rendering and analyzing the pathway to derive meaningful insights from the dataset. Starting from the KEGG pathway, it generates an interactive overlay, thus enabling interaction with selectable nodes within the KEGG pathway image. This functionality allows zooming and panning around the image, facilitating a more detailed analysis of specific pathway components. **Figure 5** shows a fraction of the Butanoate metabolism pathway.

**Figure 5:**
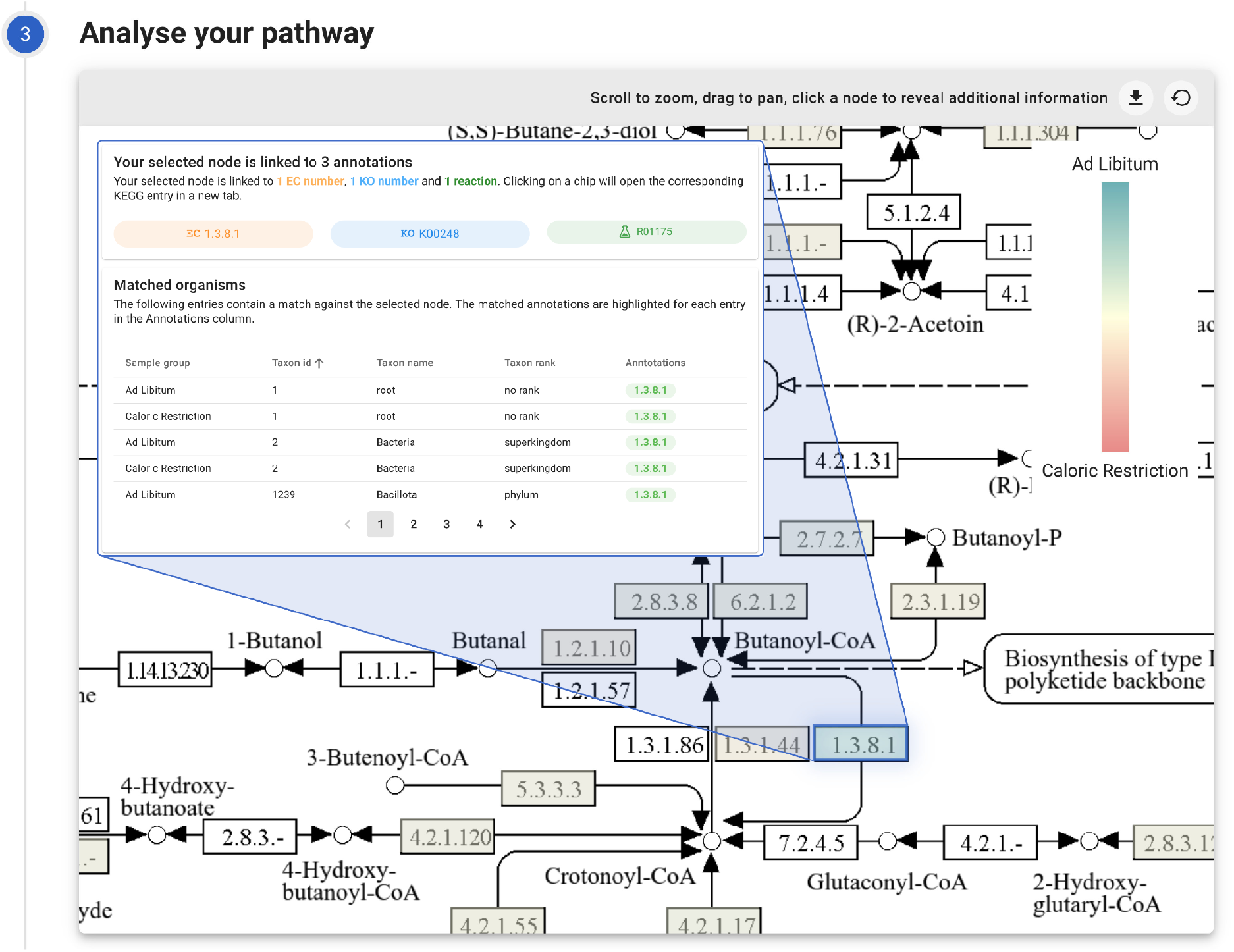
Pathway analysis. A fraction of the Butanoate metabolism pathway (map00650) is visualized. The user can analyze the pathway using the interactive pathway maps. This view shows the differential abundance for each node. Selecting a node will provide extra pathway information to the user.

Upon selection of either a compound or an enzyme/reaction, the interface is updated with additional information. Annotations linked to the chosen pathway node are listed, not only showing the evident EC numbers but also supplementary details such as reaction numbers or KEGG Ontology numbers. In addition to functional insights, taxonomic information is available as well. For each node, we ascertain a taxonomic rank and name based on matched peptides in that specific area.

Analyzing the presence in a single sample can already offer a lot of insights, but it can often be more useful to compare relationships between multiple groups or samples. One way of doing this is by looking at the difference between the two uploaded sample groups. By means of a single click, users can toggle between the different visualization views. Switching to the differential abundance view will replace the legend with a linear scale, and will show differences on top of each node using the correct color.

### 4. Exporting the results

Finally, PathwayPilot provides several options to export the results. A key export is a download of the created pathway image. Zooming and moving around the image will adjust the view, and can thus result in different exports. This way, users can select small portions of the pathway and only export the parts that are relevant for their analysis. Alongside image exports, researchers might want to further analyze results using their own tools. To make this easier, PathwayPilot provides a dense export of its internal data store. This includes mappings from input data, either peptides or proteins, to EC numbers, pathways and pathway names. This is all provided in CSV format to enable easy downstream processing and integration into other tools. Moreover, PathwayPilot also provides a pathway export, comprising an ordered list of all pathways. The ordering in this list is based on the amount of matches for that pathway, in descending order.

#### PathwayPilot in action

In order to showcase PathwayPilot, we conducted a case study using data from Tanca *et al*. [13]. This study investigated both short-term and long-term effects of caloric restriction (CR) on rat gut microbiota. They showed that a switch from *ad libitum* (AL) low fat diet to CR in young rats can induce rapid and deep changes in their gut microbiome metaproteomic profile. They observed a significant change in the expression of the microbial enzymes responsible for short-chain fatty acid biosynthesis, with CR boosting propionogenesis and limiting butyrogenesis and acetogenesis. We used the AL and CR samples and the same 32 enzymes presented in the paper as a reference for our own case study. For this case study, two out of three different study groups were extracted from the original data file. The first group contains samples from rats undergoing 3, 5 and 8 weeks of a caloric restriction (CR) diet. The second group contains samples of rats fed *ad libitum* (AL) during the same time frames of 3, 5 and 8 weeks. Then, we combined all nine AL samples, and all nine CR samples into two groups. The AL group contained a total of 80 307 peptides, while the CR group contained 70 188 peptides. The full list of peptides is available as **Additional file 1**. These two groups were then uploaded to PathwayPilot and the results were compared against the 32 reference enzymes. In order to perform the comparison, we selected a set of four different KEGG pathways that cover all the reference enzymes: (i) butanoate metabolism (map00650), (ii) carbon fixation pathways in prokaryotes (map00720), (iii) propanoate metabolism (map00640), and (iv) folate biosynthesis (map00790).

The full list of 32 enzymes, their processes as described in the original paper, EC number, and associated KEGG pathway can be found in **Table 1**. A detailed overview of the abundances is available as **Additional file 2**. The green rows in this Table indicate results that are in accordance with the original paper, the red rows indicate a different result, and the gray rows indicate EC numbers that were not reported by Unipept or do not have an associated KEGG pathway.

**Table 1.** Full list of 32 enzymes, their processes as described in the original paper, EC number, and associated KEGG pathway. Green rows indicate results that are in accordance with Tanca et al. [13], red rows indicate a different result, and gray rows indicate EC numbers that were not reported by Unipept or that do not have an associated KEGG pathway.

| Process<br>(original paper) | Enzyme | EC number | Pathway |
| --- | --- | --- | --- |
| Butyrogenesis | Acetyl-CoA acetyltransferase | 2.3.1.9 | map00650 |
| Butyrogenesis | 3-hydroxybutyryl-CoA dehydrogenase | 1.1.1.157 | map00650 |
| Butyrogenesis | Enoyl-CoA hydratase | 4.2.1.17 | map00650 |
| Butyrogenesis | Butyryl-CoA dehydrogenase | 1.3.8.1 | map00650 |
| Butyrogenesis | Butyryl-CoA:acetate CoA-transferase | 2.8.3.8 | map00650 |
| Butyrogenesis | Phosphate butyryltransferase | 2.3.1.19 | map00650 |
| Butyrogenesis | Butyrate kinase | 2.7.2.7 | map00650 |
| Butyrogenesis | Glutaconate CoA-transferase | 2.8.3.12 | map00650 |
| Butyrogenesis | (R)-2-hydroxyglutaryl-CoA dehydratase | 4.2.1.167 | None |
| Butyrogenesis | Glutaconyl-CoA decarboxylase | 7.2.4.5 | map00650 |
| Butyrogenesis | Glutamate decarboxylase | 4.1.1.15 | map00650 |
| Butyrogenesis | NAD-dependent 4-hydroxybutyrate dehydrogenase | 1.1.1.61 | map00650 |
| Butyrogenesis | 4-hydroxybutyrate coenzyme A transferase | 2.8.3.- | map00650 |
| Butyrogenesis | 4-hydroxybutyryl-CoA dehydratase/vinylacetyl-CoA-Delta-isomerase | 5.3.3.3<br>4.2.1.120 | map00650 |
| Propionogenesis | Malate dehydrogenase | 1.1.1.37 | map00720 |
| Propionogenesis | Fumarate hydratase | 4.2.1.2 | map00720 |
| Propionogenesis | Fumarate reductase flavoprotein | 1.3.1.6 | map00720 |
| Propionogenesis | Methylmalonyl-CoA mutase | 5.4.99.2 | map00720 |
| Propionogenesis | Propionyl-CoA carboxylase | 6.4.1.3 | map00720 |
| Propionogenesis | Lactoyl-CoA dehydratase | 4.2.1.54 | map00640 |
| Propionogenesis | Acryloyl-CoA reductase | 1.3.1.95 | map00640 |
| Propionogenesis | Lactaldehyde reductase | 1.1.1.77 | map00640 |
| Propionogenesis | propanediol dehydratase | 4.2.1.28 | map00640 |
| Propionogenesis | Phosphate propanoyltransferase | 2.3.1.222 | map00640 |
| Propionogenesis | Propionyl-CoA:succinate CoA transferase | 2.8.3.1 | map00640 |
| Acetogenesis | Carbon monoxide dehydrogenase | 1.2.7.4 | map00720 |
| Acetogenesis | Carbon monoxide dehydrogenase/acetyl-CoA synthase | 6.2.1.1 | map00650 |
| Acetogenesis | Formate-tetrahydrofolate ligase | 6.3.4.3 | map00720 |
| Acetogenesis | Methenyltetrahydrofolate cyclohydrolase | 3.5.4.9 | map00720 |
| Acetogenesis | Tetrahydrofolate dehydrogenase/cyclohydrolase | 1.5.1.3 | map00790 |
| Acetogenesis | Phosphate acetyltransferase | 2.3.1.8 | map00720 |
| Acetogenesis | Acetate kinase | 2.7.2.1 | map00720 |

Out of the 32 enzymes we compared, four were not reported by Unipept (Glutaconyl-CoA decarboxylase, NAD-dependent 4-hydroxybutyrate dehydrogenase, 4-hydroxybutyrate coenzyme A transferase and Propionyl-CoA:succinate CoA transferase). As no matches were found by Unipept, PathwayPilot can’t report any results for these four enzymes. There could be several reasons why Unipept did not report these peptides. Some of the peptides are semi-tryptic or non-tryptic, which are discarded by the current version of Unipept. Additionally, some peptides might not be present in the UniProtKB Swiss-Prot/TrEMBL database, which is used by Unipept, but are found in the NCBI-nr database, which was used as the search database in the original manuscript. In addition, the enzyme (R)-2-hydroxyglutaryl-CoA dehydratase is not part of any KEGG pathway and was therefore excluded from the comparison. For each of the remaining 27 enzymes, we looked at the absolute differential abundance generated by PathwayPilot. This metric shows upregulation, downregulation, or unaltered expression. This regulation can then be compared against the results from the original publication. For 24 out of 27 enzymes, we observed that the results were in accordance with the original paper. For the remaining three enzymes, we discovered a different result. Both Glutaconate CoA-transferase and Butyryl-CoA:acetate CoA-transferase showed an upregulation in the original paper, while PathwayPilot found these enzymes to be downregulated in CR samples. Lactoyl-CoA dehydratase on the other hand showed multiple fluctuations over the different samples, which indicates neither an upregulation or downregulation. PathwayPilot discovered the enzyme to be about 166% more present in CR samples. The very low number of peptides matched (24 peptides) in combination with the large number of fluctuations across samples might explain this deviation. Overall, we can see that PathwayPilot obtains results that are in accordance with those presented in the original paper, indicating its capability of creating meaningful and correct insights in the functionality of samples.

## Conclusions

We here introduce PathwayPilot, a novel tool designed to enhance the functional analysis of metaproteomic data. It stands out for its ability to provide comprehensive insights into the functional dynamics and regulatory mechanisms of microbial communities. This functionality is crucial for advancing our understanding of microbial roles in various ecosystems.

A case study has been presented to showcase the tool’s capabilities and strengths, comparing the effects of caloric restriction on the gut microbiota’s metaproteomic profile against the original results from the literature. This comparison validates PathwayPilot’s accuracy and reliability, highlighting its potential to contribute meaningful insights into microbial community research.

## Availability and requirements

Project name: PathwayPilot

Project home page: https://github.com/unipept/pathway-pilot

Operating system(s): Platform independent

Programming language: JavaScript

Other requirements: NodeJS 21 or higher

License: MIT license

Any restrictions to use by non-academics: none

## List of abbreviations

EC: Enzyme Commission
GO: Gene Ontology
KEGG: Kyoto Encyclopedia of Genes and Genomes
CR: Caloric Restriction
AL: Ad Libitum

## Declarations

### Ethics approval and consent to participate

Not applicable

### Consent for publication

Not applicable

### Availability of data and material

All data generated or analyzed during this study are included in this published article and its supplementary information files

### Authors’ contributions

TVM has made substantial contributions to the design of the work and the creation of new software used in the work. TVM and TVDB have drafted the work. All authors have made substantial contributions to the conception of the work, have revised the work and approve the submitted version, and agree to be personally accountable for their own contributions and to ensure that questions related to the accuracy or integrity of any part of the work, even ones in which the author was not personally involved, are appropriately investigated, resolved, and the resolution documented in the literature.

## Acknowledgements

The development of this tool was initiated at the EuBIC-MS Developers’ meeting in Switzerland [14]. This work has benefited from collaborations facilitated by the Metaproteomics Initiative (https://metaproteomics.org/) whose goals are to promote, improve, and standardize metaproteomics [15].

